# Fluorometric intracellular Na^+^ measurement as compared with flame emission assay: An unexpected problem with gramicidin-based calibration

**DOI:** 10.1101/713792

**Authors:** Valentina E. Yurinskaya, Nikolay D. Aksenov, Alexey V. Moshkov, Tatyana S. Goryachaya, Alexey A. Vereninov

## Abstract

Fluorescent probes are a popular and indispensable tool for monitoring sodium concentration in living cells in situ. Calibration of fluorescent probes inside cells commonly uses ionophores to equilibrate intracellular and external ion concentrations. Here we test this calibration method using in parallel classical flame emission assay. Suspension human lymphoma cells allow both flow cytometry fluorometric study and flame emission assay. The most sensitive Na^+^ fluorescent probe ANG-2 and the most common ionophores were tested. Cellular Na^+^ was altered for calibration in three different ways: by stopping the sodium pump with ouabain, by inducing of apoptosis with staurosporine, and by gramicidin or amphotericin B treatment. We found that ANG-2 fluorescence in cells treated with gramicidin or amphotericin was about two fold lower than in the cells with the same sodium concentration but without ionophores. The equal fluorescence measured in the absence and in the presence of ionophores corresponds to different cell sodium concentrations. No effect of gramicidin on hydrolyzed ANG was observed *in vitro*. The mechanism, by which gramicidin decreases ANG fluorescence in cells is unlikely to be physical quenching and remains obscure. We conclude that ANG fluorescence does not display realistic cell Na^+^ if fluorescence in cell is measured in ionophore absence while calibrated in its presence.

## Introduction

Monovalent ions underlie fundamental cell functions: water balance and electric processes, intra- and intercellular signaling, cell movement, pH regulation and metabolite transport into and out of cells. Optical methods of monitoring intracellular monovalent ions are very important because of the complexity of multicellular organs and tissues and lability of ions in living cells. Using ion-sensitive optical probes in flow cytometry allows combining precise single cell photometry with analysis of cell diversity in large populations. Calibration of optical signals is an important problem when cell ion content is measured by optical methods. It is based on the use of ionophores to equilibrate intracellular and external ion concentrations in cells. Although X-ray elemental analysis allows, in principle, to measure intracellular ion concentrations with high spatial resolution, it is very laborious and not applicable to living cells. We aimed to compare data on cell Na^+^ obtained in human U937 lymphoma cells by flame emission assay and by flow cytometry using the Na^+^-sensitive probe Asante Natrium Green-2 (ANG). The other new feature of our study is that cell Na^+^ was altered not only by ionophores but also by stopping the sodium pump with ouabain or by inducing apoptosis with staurosporine (STS). Alteration of monovalent ion balance in U937 cells in these cases has earlier been studied by us in detail (Yurinskaya et al., 2005*a*, 2010, 2011; Vereninov et al., 2007, 2014, 2016). We report here that the same intracellular Na^+^ in ionophore-treated and untreated cells (as reported by flame emission analysis) produces different ANG fluorescence indicating that gramicidin reduces ANG fluorescence. We conclude that ANG fluorescence calibrated with ionophores does not display realistic cell Na^+^ if fluorescence is measured in ionophore absence while calibrated in its presence. ANG fluorescence decrease in cells due to ionophores, gramicidin (Gram) in particular, does not preclude their using for monitoring relative changes in intracellular Na but much precaution is required for quantitative [Na^+^]_in_ determination.

## Methods

### Cell culture and treatment

The human histiocytic lymphoma cell line U937 was obtained from the Russian Cell Culture Collection (Institute of Cytology, Russian Academy of Sciences, cat. number 220B1). Cells were cultured in RPMI 1640 medium (Biolot, Russia) supplemented with 10% fetal bovine serum (HyClone Standard, USA) at 37 °C and 5 % CO_2_. Cells were grown to a density of 1 × 10^6^ cells per ml and treated with 10 µM ouabain (Sigma-Aldrich) or with 1 µM STS (Sigma-Aldrich) for indicated time. Gramicidin (from Bacillus aneurinolyticus) and amphotericin B (AmB) were from Sigma-Aldrich and prepared as stock solutions in DMSO at 1 mM and 3 mM, respectively. Na^+^-specific fluorescent dye Asante Natrium Green-2 AM ester (ANG-AM) was from Teflabs (Austin, TX, Cat. No. 3512), distributed by Abcam under the name ION NaTRIUM Green™-2 AM (ab142802). 5 µM Gram was added for 30 min simultaneously with ANG, 15 µM AmB was added during the last 15 min of a 30-min ANG loading.

### Flow cytometry and cell Na^+^ content determination

A stock solution of 2.5 mM ANG-AM was prepared in DMSO +10% Pluronic (50 µg ANG-AM were dissolved in 10 µL DMSO and mixed 1:1 with 20% (w/v) Pluronic F-127 stock solution in DMSO). After intermediate dilution in PBS, ANG-AM was added directly into the media with cultured cells to a final concentration of 0.5 µM. Incubation of cells with ANG-AM was carried out for 30 min at room temperature (∼23°C). Stained cells were analyzed on a CytoFLEX Flow Cytometer (Beckman Coulter, Inc., CA, USA). ANG fluorescence was excited using a 488 nm laser, emission was detected in PE (PhycoErythrin) channel with a 585/42 nm bandpass filter (marked below in figures as ANG, PE). All fluorescence histograms were obtained at the same cytometer settings for a minimum of 10,000 cells. Cell population P1 was gated by FSC/SSC (forward scatter/side scatter) with a threshold set at 1×10^4^ to exclude most of microparticles and debris (Yurinskaya et al., 2017 (**Figure 1A**). The area (A) of flow cytometer parameters (FSC-A, SSC-A, PE-A) was used.

**Figure 1.**
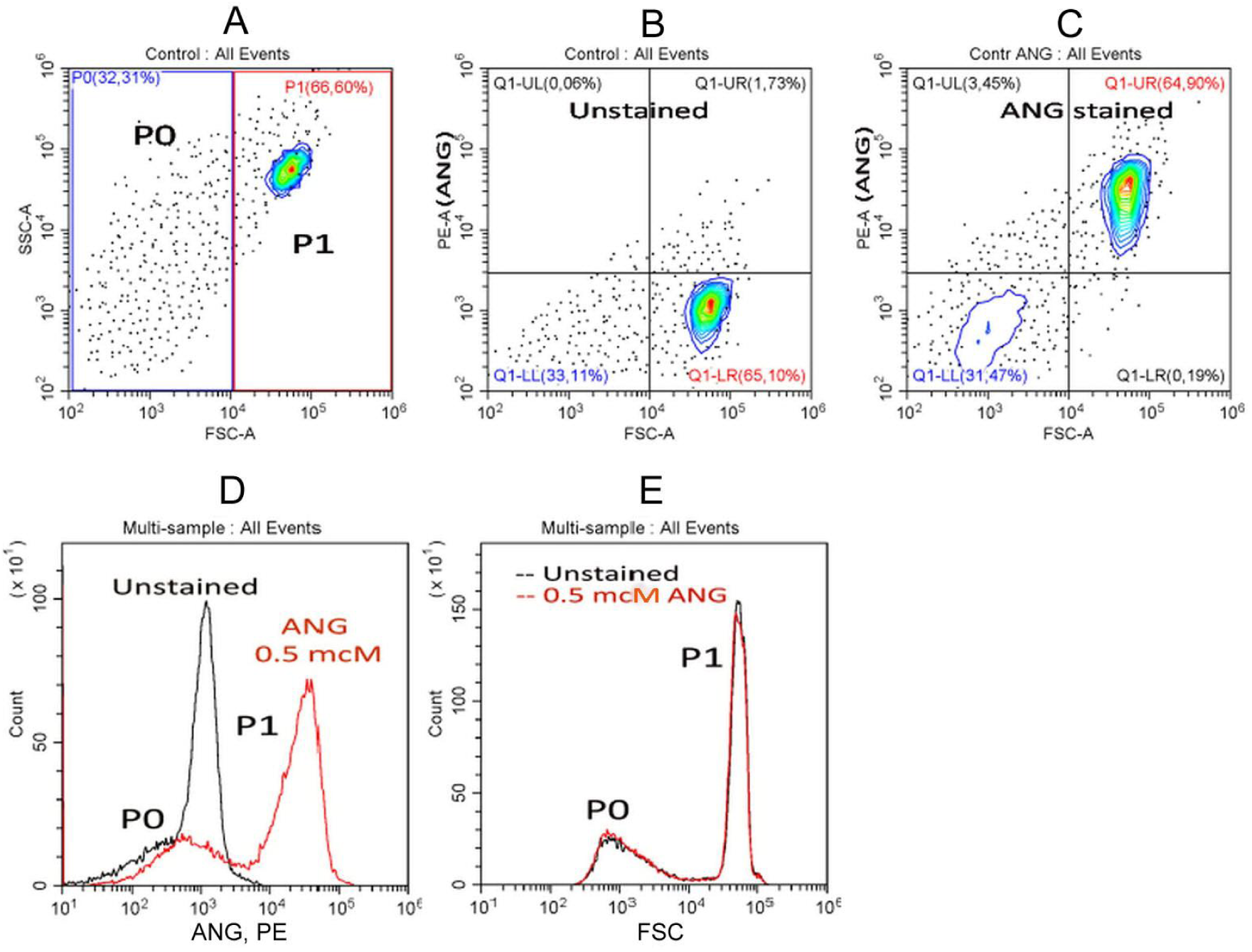
ANG fluorescence (PE), side scatter (SSC) and forward scatter (FSC) distributions of normal U937 cells in unstained (A, B) and ANG-stained (C) total cell cultures. Staining was achieved by exposing cells to 0.5 µM ANG-AM for 30 min, as described in Methods. The displayed data were all obtained on the same day and represent at least eight separate experiments.

Cell Na^+^ content was determined by flame emission on a Perkin-Elmer AA 306 spectrophotometer as described in detail earlier (Yurinskaya et al. 2005*a*, *b*, 2011; Vereninov et al. 2007, 2008) and evaluated in mmol per g of cell protein, Na^+^ concentration is given in mM per cell water. Cell water changed insignificantly under all tested conditions and was 5.8 ml per g of cell protein on average (as measured by buoyant density, not shown). Cell volume was measured additionally by Scepter cell counter equipped with a 40-μm sensor and software version 2.1, 2011 (Merck Millipore, Germany) as described earlier (Yurinskaya et al., 2017).

### Fluorescence microscopy

Images of ANG-stained cells were acquired on Leica TCS SP5 confocal microscope. Cells were placed on a cover-slip in a plexiglass holder and excited at 488 nm through a 40x Plano oil-immersion objective; fluorescence emission was collected at 500 – 654 nm. Laser power, rather than PMT voltage, was varied to avoid image saturation for the observation of control and treated cells.

Fluorescence of hydrolyzed ANG in water solution was measured using Terasaki multiwell plate, using a 30x water-immersion objective and a microscope equipped with epi-fluorescent attachment. Fluorescence was excited by LED (5 W, 460 nm, TDS Lighting Co) and passed through dichroic filter (480-700 nm). A charged form of ANG was obtained by hydrolysis according to the recommended protocol (Molecular Probes Handbook, 2010). The effect of Gram, Na^+^, and K^+^ on hydrolyzed ANG fluorescence was tested by adding 1.5 µL water solution of Gram (final 8 µM) with DMSO (final 0.08%), or DMSO (0.08%), or of H2O to 16.5 µL of solution in the plate well contained ANG 55 µM, KCl 125 mM, NaCl 40 mM, pH 7.0.

### Statistical and data analysis

CytExpert 2.0 Beckman Coulter software was used for data analysis. The means of PE-A for appropriate cell subpopulations given by CytExpert software were averaged for all samples analyzed. Data were expressed as mean ± SD for indicated numbers of experiments and analyzed using Student’s t test. P < 0.05 was considered as statistically significant.

## Results

### P1 and P0 subsets in ANG and FSC flow diagrams of the total U937 population

Flow cytometric analysis of the original U937 cell culture shows two main particle subsets in FSC/SSC and FSC/PE(ANG) plots that we denote as P0 and P1 (**Figure 1A-C**). P1 represents intact cells while P0 contains particles, vesicles or debris (Yurinskaya et al., 2017). Cell culture as a whole is analyzed using the method of flame emission, and it is usually unknown which part consists of cells, and which of the fragments of cells and debris. Flow cytofluorometry with ANG allows to answer this question. Fluorescence intensity plots contain the P0 and P1 subsets similar to those in FSC histograms, but with slightly broader peaks (**Figure 1D, E**). Comparison of **Figures 1B** and **1C** show that ANG fluorescence (PE-A signal) increases noticeably for all P1 cells and only slightly for P0 subset. Flow cytometry of ANG-stained U937 cells allows evaluation of uniformity of cell populations used in the flame emission assays. The cumulative ANG signal associated with P1 (calculated from the mean fluorescence and the number of particles) accounts for about 98% of all the ANG contained in P1+ P0. The typical P0/P1 ratio for the integral ANG signal was 0.019±0.001 (n = 8). A similar ratio for the FSC signal was 0.023±0.002. That means that sodium reported by the flame emission analysis practically corresponds to the P1 subset. Therefore, we further consider only ANG fluorescence of the P1 subset. Microscopy confirms the uniform distribution of ANG staining in P1 cells and a more or less homogeneous ANG distribution inside the cells (**Figure 2**). It should be noted that flow cytofluorometry has a definite advantage compared with quantitative fluorescence microscopy of ANG-Na due to more accurate optical measurements and high statistics (10-20 thousand cells).

**Figure 2.**
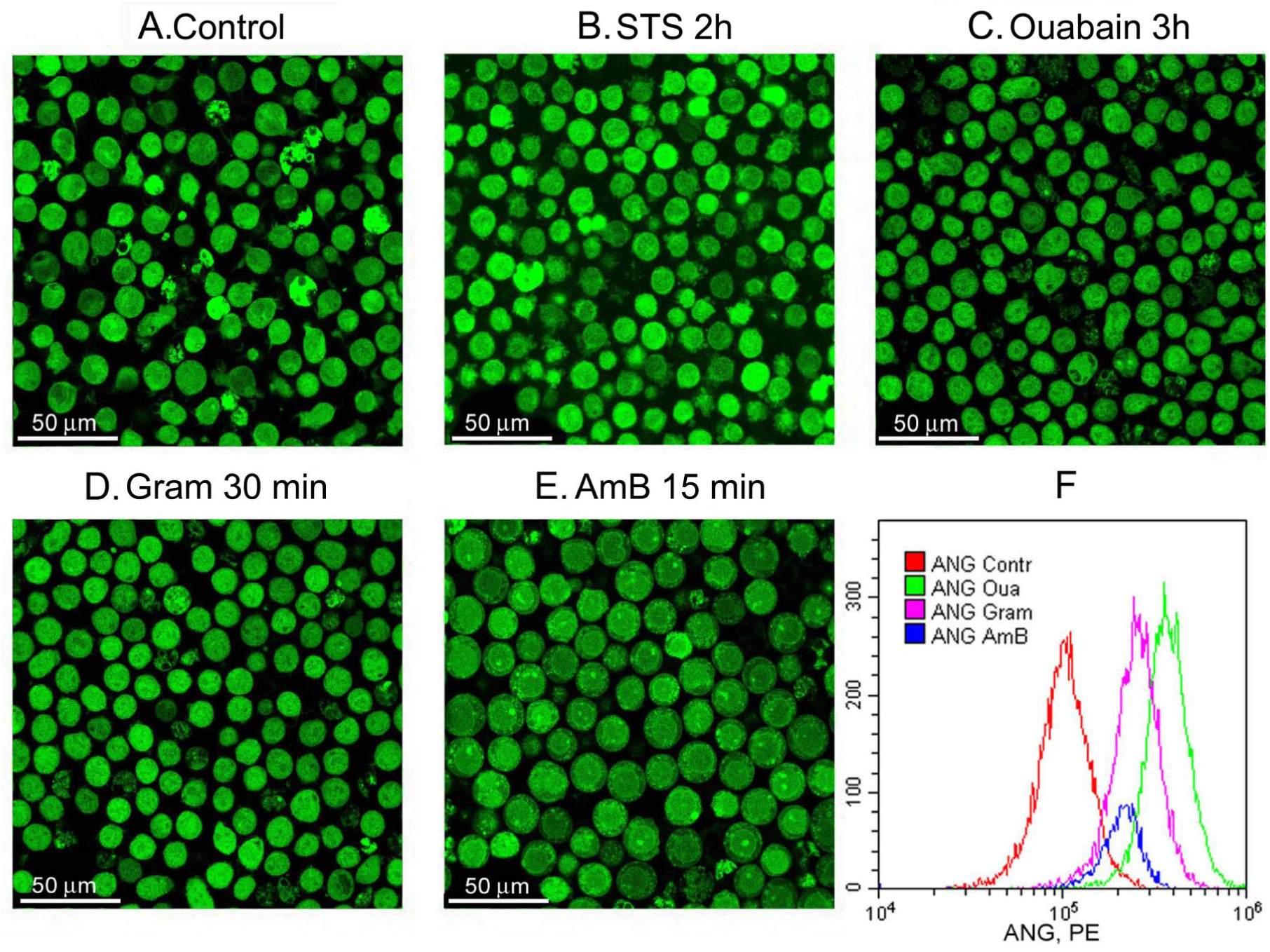
Microscopy of ANG stained cells treated with STS, ouabain, Gram or AmB. U937 cells were treated with STS **(B)** or ouabain **(C)** during indicated time and stained with ANG-AM for last 30 min as described in Methods. Alternatively, cells without treatment with ouabain or STS were stained with ANG-AM in presence of Gram **(D)** or AmB **(E)** for the last 15 min of 30-min ANG-AM staining. After staining cells were analyzed on a flow cytometer **(F)** or viewed on a confocal microscope **(A-E)**. Fluorescence images were acquired on the microscope at a laser power of 100% for control and for STS-treated cells, 30% for ouabain-treated cells, 40% for gramicidin-treated cells and 70% for AmB treated cells.

Cell loading has been tested at several concentrations of ANG (**Figure 3A**). The concentration of 0.5 µM was chosen as sufficient because ANG fluorescence of the P1 subset significantly overlaps the background cell autofluorescence (**Figure 1D, 3D**). The protocol recommended by the manufacturer calls for loading of ANG for 30 min followed by a wash step and additional incubation, to allow complete deesterification of the dye. Such a procedure has often been used to prepare cells for microscopic observation, but we found that washing of cells to remove the external ANG had no significant impact on the ANG signal measured on a flow cytometer (**Figure 3C)**. Despite the fact that the presence of serum in the loading medium decreased cell fluorescence, the signal remained sufficiently strong (**Figure 3C, 3D**), and we preferred to keep the serum. Thus, a 30 min incubation with 0.5 µM ANG in normal media with serum without a subsequent wash have been used in all experiments. Importantly, this staining procedure did not affect the FSC/SSC histogram, which is known to be a sensitive indicator of cell health (**Figure 1E**).

**Figure 3.**
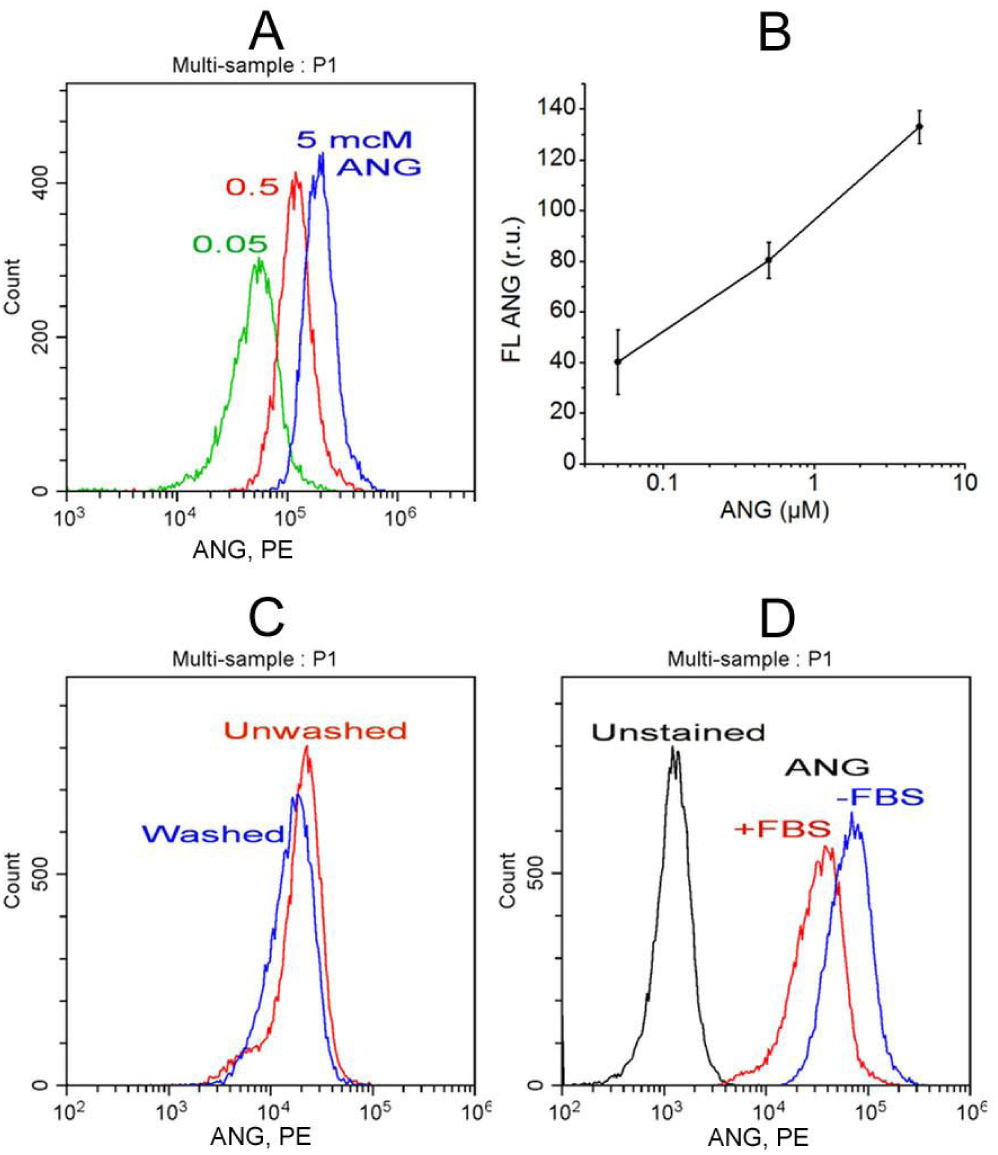
Effects of ANG concentration (A, B), washing (C) and serum (D, FBS) on ANG fluorescence in the P1 cell subset. The displayed histograms were obtained in independent experiments at identical instrument settings. (**B**) dependence of cell ANG fluorescence on ANG-AM concentration in the serum free RPMI media, means ± SD for 2 experiments like in **Figure 3A**.

### Comparison between [Na^+^]_in_ estimation by ANG fluorescence and by flame emission assay

Ionophores are commonly used to calibrate fluorescent ion probes by controlling intracellular ion concentration. In this study, we did not need the usual step of calibration and applied a different approach. We compared the relative changes in cell sodium, as measured by flame emission analysis, with the relative changes in ANG-Na fluorescence detected on a flow cytometer, which reflects the total fluorescence of individual cells, averaged over 20 thousand measurements. Increase in cell Na^+^ was induced by stopping the pump with ouabain or by STS; the effects of these stimuli on cell ion composition have been previously characterized in detail (Yurinskaya et al., 2005, 2010, 2011, 2017, 2019; Vereninov et al., 2007, 2014, 2016). The treatment of cells with Gram commonly used in ANG calibration gave another way to increase cell Na^+^. Indeed, both ouabain and Gram increased ANG fluorescence significantly (**Figure 4A, 4B**). Cell volume measured by Scepter coulter practically did not change in cells treated with ouabain or gramicidin for the indicated time (**Figure 5**). FSC histograms do not change in these cases as well (**Figure 4C, 4D**). FSC is usually considered as an indicator of cell size, albeit with some reservations (Shapiro, 2003; Tzur et al., 2011; Yurinskaya et al., 2017).

**Figure 4.**
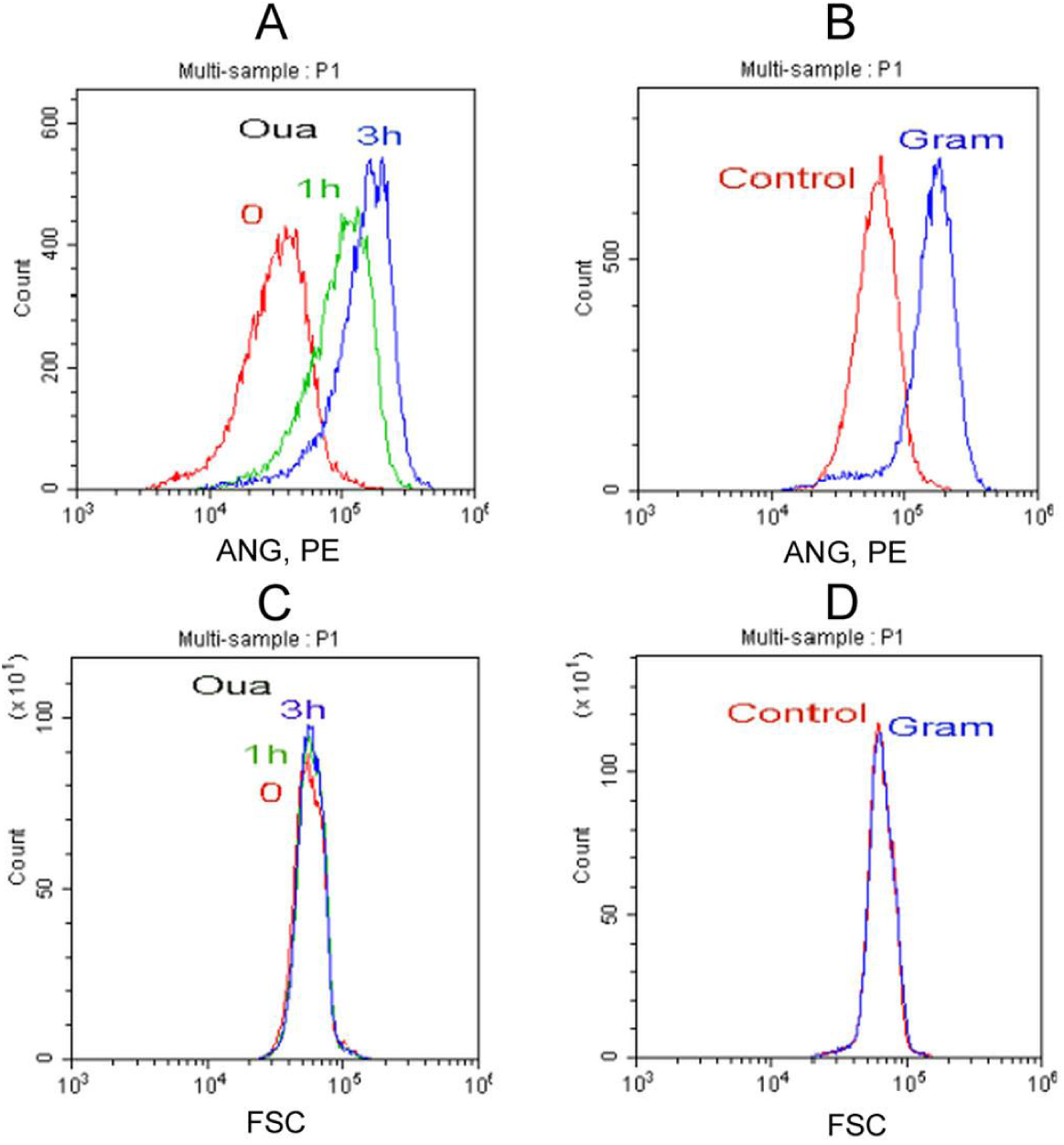
ANG fluorescence- (A, B) and FSC-histograms (C, D) for the ouabain and gramicidin treated cells. Histograms were obtained in independent representative experiments at the same instrument settings for the compared data. Cells were treated with 10 µM ouabain for indicated time; 5 µM gramicidin was added for 30 min with ANG-AM.

**Figure 5.**
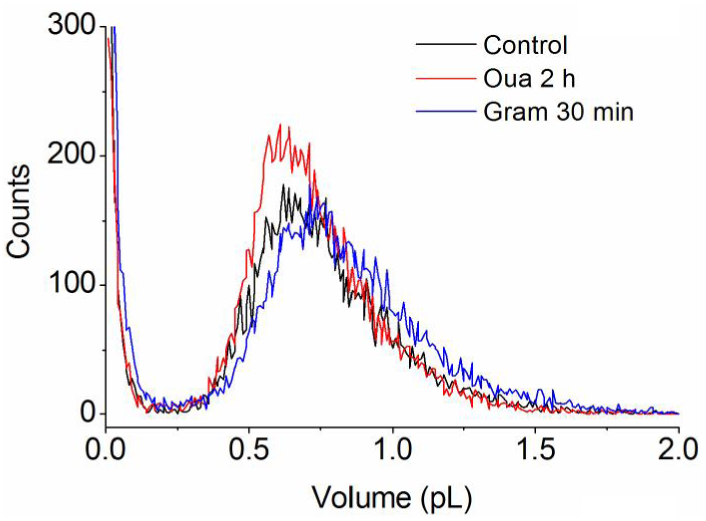
Cell volume histograms obtained by Scepter coulter for cell cultures treated with 10 µM ouabain or 5 µM Gram for indicated time. Compared histograms are obtained in the same representative experiment.

Flow cytometry showed an excellent agreement of changes in fluorescence and in cell Na^+^ increase due to stopping the pump with ouabain and STS-induced apoptosis (**Figure 6A, 6B)**. Gram and AmB added to U937 cells increased intracellular Na^+^ to a greater extent than did ouabain as obtained by flame emission assay. However, the increase in ANG fluorescence was much smaller (**Figure 6A**). The discrepancy between the fluorescence of ANG and Na^+^ concentration by flame emission in the presence of ionophores assay is especially striking when comparing the data on the same graph (**Figure 6C**). It should be stressed that all compared flow cytometry data were obtained at the same instrumental settings and were well reproduced. We conclude that ANG fluorescence does not display realistic cell Na^+^ if fluorescence is measured in ionophore absence while calibrated in its presence.

**Figure 6.**
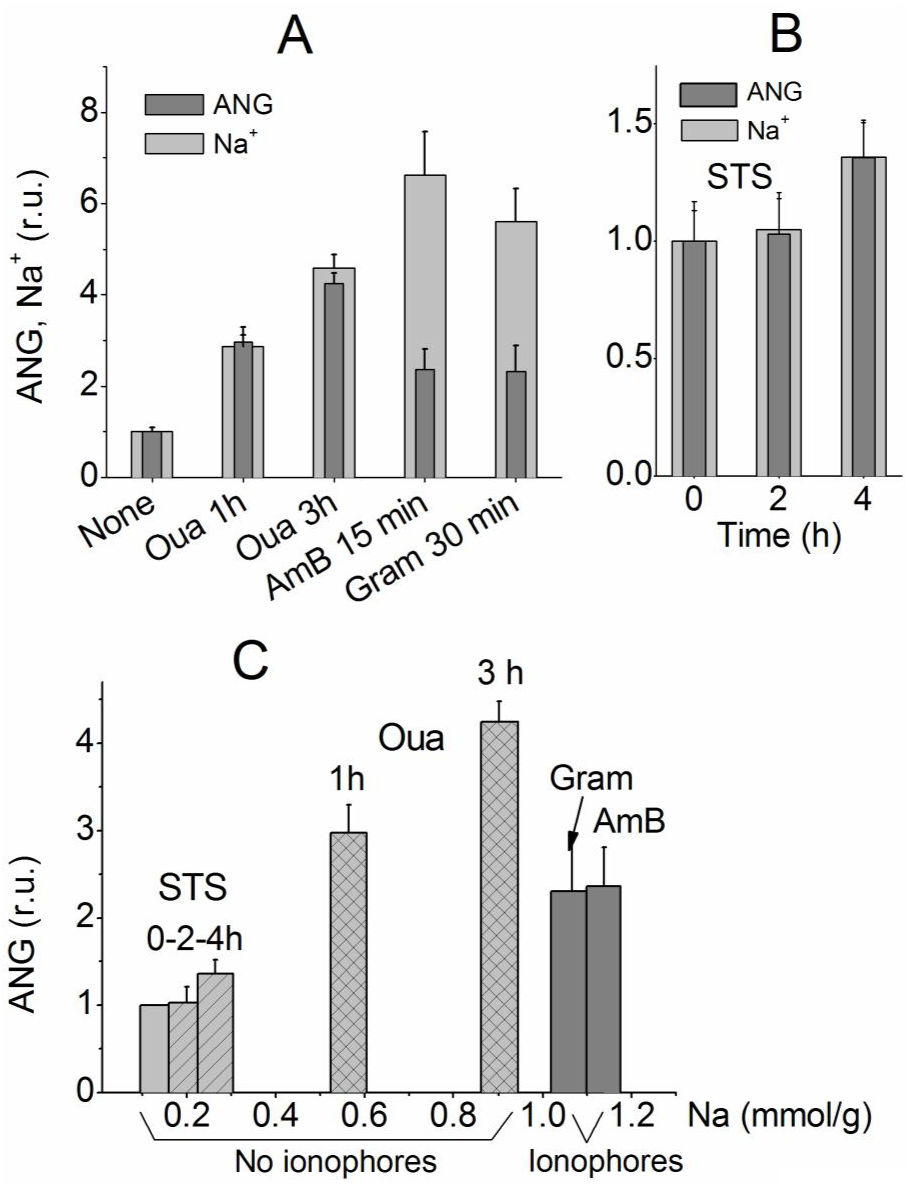
Comparison of changes in ANG fluorescence and in cellular Na^+^ by flame emission assay under various conditions. (A) cells were treated with 10 µM ouabain or 5 µM Gram, or 15 µM AmB for indicated time; (B) cells were treated with 1 µM STS inducing apoptosis; (C) ANG fluorescence at different Na^+^ obtained with and without ionophores. Other indications are on the graphs. Means ± SD were normalized to the values in untreated cells for 3-5 experiments (in duplicate for cell Na^+^ content). The measured Na^+^ content in mmol/g cell protein corresponds to intracellular concentrations ranging from 34 to 154 mM.

### Gramicidin does not influence ANG fluorescence *in vitro*

We tested the effect of Gram on hydrolyzed ANG. ANG fluorescence was measured using Terasaki multiwell plate and fluorescent microscope as described in Methods. The ANG-AM used for probing intracellular Na^+^ is practically nonfluorescent *in vitro* (**Figure 7**). It is believed that after penetrating the cell membrane ANG-AM is hydrolyzed by non-specific esterases. Indeed, after hydrolysis ANG-AM *in vitro* by known protocol (Molecular Probes Handbook, 2010) ANG strongly responds to Na^+^ but relatively weekly to K^+^ supporting the view that the same can occur in cells (**Figure 7**). The most important finding with regard to our problem was the lack of any quenching effect of Gram on fluorescence of hydrolyzed ANG at ion concentrations imitating those in the cytoplasm.

**Figure 7.**
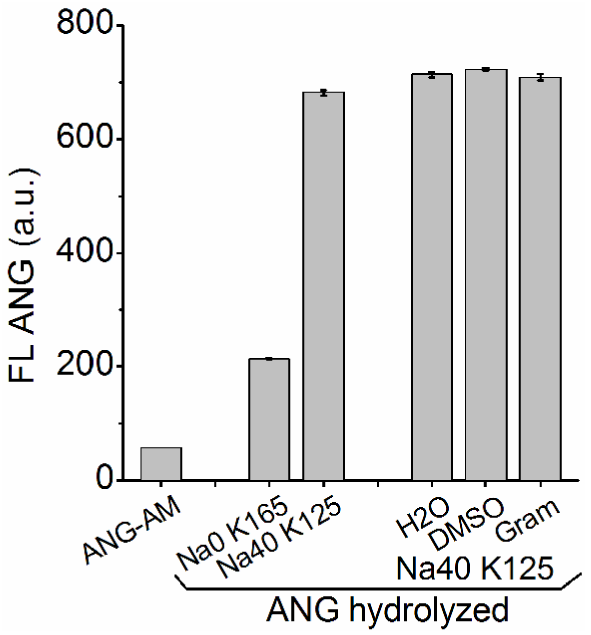
The effect of Na^+^, K^+^ and Gram on fluorescence of ANG-AM and on the hydrolyzed ANG in water solutions. Averages for 2 measurements.

## Discussion

We showed, first, that all the cells assayed by flame emission and by flow cytometry make a more or less homogeneous group. Then we applied flow cytometry to compare two different methods of Na^+^ measurement. To our knowledge, this is the first report of using ANG in flow cytometry and the first attempt to compare Na^+^ data obtained with optical probes and by direct flame emission assay. The most important advantage of flow cytofluorometry compared to quantitative ANG-Na fluorescence microscopy is that the flow cytometer determines the total fluorescence of an individual cell, and there is no uncertainty as to whether the concentration or amount of ANG-Na is in the area inside the cell selected for photometry. ANG and ion-sensitive probes in cuvette reflect (due to change in fluorescence intensity or spectra) ion concentration. In the case of a cell, the fluorescence intensity of ANG-Na in the selected area of image should reflect the local concentration of ANG-Na. The total amount of ANG-Na in the whole cell in this case depends on the size-volume of the cell. To compare flame emission data with optical data, the cell volume must be known in addition to local fluorescence. That is always problem.

ANG-2 is currently the most popular probe to study sodium distribution and dynamics in various types of cells. ANG has been used in brain slices, neurons, their dendrites and axons, astrocytes (Lamy and Chatton, 2011; Schreiner and Rose, 2012; Sarkar et al., 2014; Miyazaki, Ross, 2015; Rose and Verkhratsky, 2016; Breslin et al., 2018; Noor et al., 2018), HEK293 cells (Mahmmoud et al., 2014), cockroach salivary gland cells (Roder, Hille, 2014), cardiomyocytes (Kornyeyev et al., 2016), and prostate cancer cell lines (Iamshanova et al., 2016). Optical Na^+^ monitoring is quite indispensable in neuroscience because of the extreme complexity of the object structure. The number of studies using fluorometric Na^+^ measurements is rapidly increasing (Gao et al., 2017).

The use of ionophores is the main approach to convert fluorescence into absolute values of Na^+^ concentration. In this study we compared data on intracellular Na^+^ in human U937 lymphoma cells obtained by flow cytofluorometry using ANG and by the well-established flame emission assay. Cellular Na^+^ was altered in three different ways: by stopping the sodium pump with ouabain, by inducing apoptosis with staurosporine, and by treating cells with Gram. We found that ANG fluorescence in cells treated with gramicidin or amphotericin was about two fold lower than in the cells with the same sodium concentration but without ionophores.

Our tests of interactions of hydrolyzed ANG with Gram *in vitro* lead to a conclusion that a decrease in ANG fluorescence in cells caused by Gram is hardly due to “simple” physical quenching of ANG in cell. It is known that complex ANG with Na^+^ in cells differ from the analogous complex in water solution by K_d_ (Lamy and Chatton, 2011; Iamshanova et al., 2016). Similar differences were found earlier for other ion-specific probes, e.g. for Sodium Green (Amorino, Fox, 1995). Mechanism of this effect remains unknown. It has been shown for water solutions that K^+^ can change the dependence of the sodium probe fluorescence on Na^+^ concentration (see e.g. Iamshanova et al., 2016; Naumann et al., 2018). Harootunian with colleagues in their pioneering work (1989) supposed that SBFI spectral characteristics in cytoplasm could be different in cell and in external water media due to differences in viscosity. Other more general and speculative hypotheses could be that an influence of gramicidin on ANG fluorescence in cell is determined by changes in interaction of ANG with some unknown cytoplasm components or an influence of the process of “de-estherification” of the ANG-AM in cells. It is for this reason the calibration *in situ* is always performed.

In view of the gramicidin effect on ANG fluorescence observed in our experiments, the calibration without ionophores attracts attention. Rose with colleagues compared traditional calibration using gramicidin with calibration without gramicidin by direct dializing of single cell via patch pipette by saline with different Na^+^ (Mondragão et al., 2016). They believed that similar results were obtained. It should be noted that these authors study changes in intracellular Na^+^ using SBFI in the ratiometric mode.

An interesting question concerns equalizing extra- and intra-cellular concentrations of cations in cells treated with Gram and other ionophores. Our flame emission assay showed that there was no exact equalization of monovalent cation concentrations in the medium and cells after their treatment with Gram. Na^+^ level becomes somewhat higher than in the medium, while K^+^ level is about 25 mM at external concentration 5 mM after 30 min treatment of U937 cells with 5 µM gramicidin (**Figure 8**).

**Figure 8.**
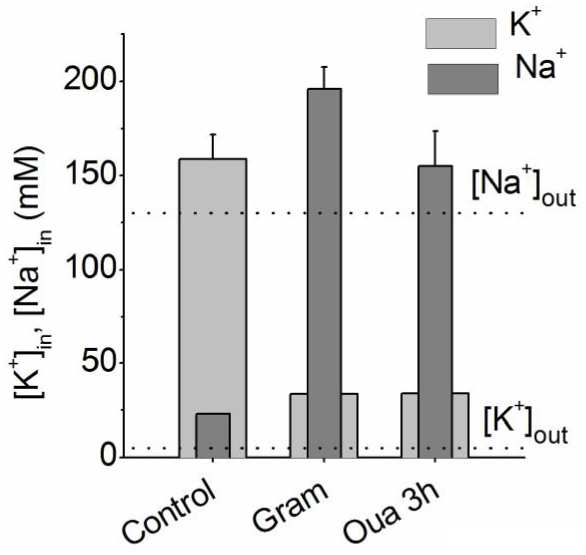
[K^+^]_in_ and [Na^+^]_in_ changes in U937 cells treated with 5 µM Gram for 30 min or with 10 µM ouabain (Oua) for 3 h. Data obtained by flame emission assay. Mean ± SD from 3 experiments with duplicate determinations. Dot lines indicate [Na^+^]_out_, [K^+^]_out_

It is believed that in the media where Cl^−^ is partly replaced with gluconate the Donnan’s rules become insignificant (Harootunian et al., 1989). We calculated changes in K^+^ and Na^+^ concentration using our recent computational tool based on the Donnan’s rules (Yurinskaya et al., 2019) for model cell with parameters similar to U937 cells. It appeared that at increasing of the cell membrane permeability for K^+^ and Na^+^ (pK, pNa in calculation) by about 10 and 50 times, imitating the gramicidin effect, the K^+^ level in cell is decreased, but remains higher than extracellular level, while Na^+^ is increased slightly higher than extracellular (**Figure 9**).

**Figure 9.**
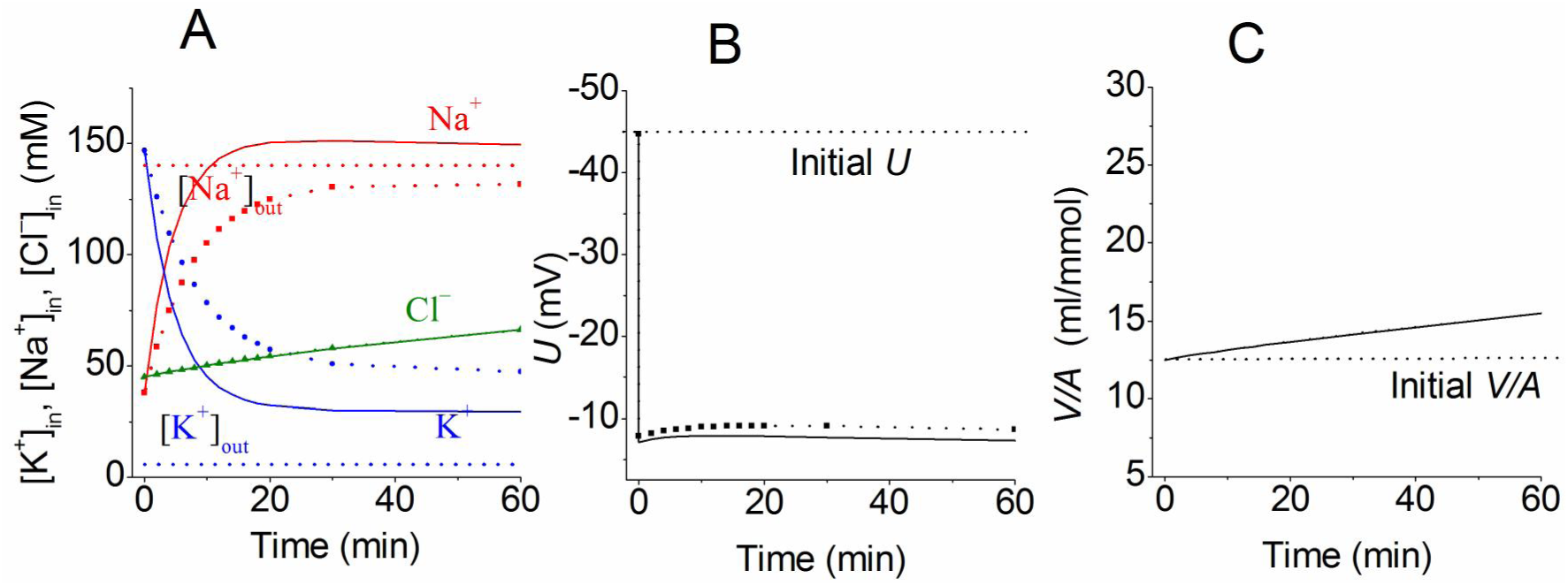
[K^+^]_in_, [Na^+^]_in_, and [Cl^−^]_in_, membrane potential (*U*) and cell volume (*V/A*) calculated for model cell with parameters similar to U937 cells. In balance state (t<0), in mM [Na^+^]_in_ 38, [K^+^]_in_ 147, [Cl^−^]_in_ 45, V/A 12.50 pNa0.00382 pK0.022 pCl 0.0091; Gram is added at t=0 and pNa, pK are set 0.1 (dot lines with symbols) or 0.2 (solid lines). Other parameters remain unchanged. Dash lines indicate [Na^+^]_out_, [K^+^]_out_ (**A**) and initial parameters on (**B, C**). Cell volume is given in ml per mmol of impermeant intracellular anions (*V/A*).

Calculation showed that chloride permeability of the cell membrane is ordinarily the most important determinant of the rate of the monovalent ion redistribution after blocking the Na/K pump or an increase in Na^+^ and K^+^ permeabilities due to ionophores. It is for this reason a rather long time is required for the “standard” cell like U937 to swell after treatment with ouabain or Gram and much less time after treatment with amphotericin which increases not only Na^+^ and K^+^ but also Cl^−^ permeability of the cell membrane.

Our considerations on equalizing intra- and extra-cellular Na^+^ concentrations are consonant with the opinion of Boron with colleagues, who were among the few who performed flame emission analysis in parallel with determination of intra-cellular Na^+^ in HeLa cells using SBFI probe and various ionophores. They note that ionophores (gramicidin, nigericin, monensin) do not solve the problem equalization of Na^+^ concentration across the cell membrane (Zahler et al., 1997).

### Our general conclusion

A decrease in ANG fluorescence due to ionophores, gramicidin in particular, does not preclude its use for monitoring relative changes in intra-cellular Na^+^, but much precaution is required for quantitative [Na^+^]_in_ determination.

## Acknowledgments

The authors are grateful to Dr. Michael Model (Department of Biological Sciences, Kent State University, Kent, Ohio 44242, USA) for manuscript revision and suggesting improvements. The research was supported by the Grants of Russian Federation (No. 0124-2019-0002, No. 0124-2019-0003). The authors declare that the research was conducted in the absence of any commercial or financial relationships.

## Author contributions

All authors contributed to the design of the experiments, performed the experiments, and analysed the data. AV wrote the manuscript with input from all authors. All authors have approved the final version of the manuscript and agree to be accountable for all aspects of the work. All persons designated as authors qualify for authorship, and all those who qualify for authorship are listed.

